# scXpand: Pan-cancer detection of T-cell clonal expansion from single-cell RNA sequencing without paired single-cell TCR sequencing

**DOI:** 10.1101/2025.09.14.676069

**Authors:** Ofir Shorer, Ron Amit, Keren Yizhak

## Abstract

Advances in single-cell sequencing have enabled detailed characterization of T-cell clonal dynamics in cancer. However, analyses aiming to link transcriptional landscape to T-cell clonality remain limited by confounding factors unequally controlled in different studies. To address this challenge, we developed scXpand, the first machine learning framework for pan-cancer detection of T-cell clonal expansion directly from single-cell RNA sequencing (scRNA-seq), without paired T-cell receptor (TCR) sequencing. Trained and tested using our in-house-constructed human pan-cancer database of paired scRNA/TCR-seq profiles from 2.6 million T cells, scXpand demonstrates robust and accurate detection of clonal expansion across tissues and T-cell subtypes. Applied to datasets lacking TCR sequencing, scXpand predictions correspond with known characteristics of the tumor microenvironment. Overall, scXpand is the first framework to detect T-cell clonal expansion across cancers directly from scRNA-seq, enabling broad use on datasets lacking scTCR-seq, while supporting scalable, memory-efficient processing, including pre-trained models with user-friendly documentation for flexible applications.

## Introduction

The recent advances of sequencing at the single-cell level enabled us to characterize the cellular landscape of diverse tissues across various pathological conditions, such as cancer. More specifically, single-cell RNA sequencing (scRNA-seq) and single-cell T-cell receptor (TCR) sequencing (scTCR-seq), provided us with better understanding of T-cell immune responses and T-cell clonal dynamics in the context of cancer therapy and clinical outcomes^1–4^. In turn, identification of expanded T-cell clones and their phenotypic landscape, as well as the source of the intra-tumoral TCR repertoire, were previously linked to immunotherapy outcomes across multiple cancer types^5–8^. However, despite their unprecedented potential of linking transcriptional landscape to T-cell clonality, such analyses of paired scRNA/TCR-seq data are still highly limited by confounding and technical factors unequally controlled in different studies^1^. These include different scopes of TCR capture due to the varying size and location of each tumor sample, as well as sample quality and limited number of productive TCRs following sequencing. As a result, when observing a single T cell having a unique TCR sequence not shared with any other T cell (termed ‘singleton’), it is uncertain whether such singleton represents a larger T-cell clone masked by these confounding factors. Such a discrepancy can potentially be addressed using the transcriptional phenotype of T cells, enabling the detection of those that are members of expanded clones, even without the availability of TCR sequencing data.

Multiple studies developed computational tools utilizing paired scRNA/TCR-seq datasets, addressing specific aspects of the data to provide additional information for downstream analysis. For example, Zhu et al. developed a deep learning method (scNAT) for integration of paired scRNA/TCR-seq data and its representation using a latent space for further analysis^9^. Schattgen et al. introduced clonotype neighbor graph analysis (CoNGA), which identifies correlations between gene expression data and TCR sequencing to elucidate complex relationships between TCR sequence and T-cell phenotype^10^. Zhang et al. developed ‘tessa’, a tool for integration of scTCR-seq and gene expression, resulting with TCR networks that are reflective of T-cell functional status^11^. More recently, Tan et al. introduced ‘predicTCR’, a machine learning classifier trained on scRNA/TCR-seq data, which identifies individual tumor-reactive T cells based on scRNA-seq data alone^12^. However, there is currently no method utilizing single-cell data of human cancer patients for the prediction of T-cell clonal expansion at the pan-cancer level, spanning multiple tissues and subtypes of T cells.

To address this challenge, we developed scXpand, a machine learning framework trained and tested using our in-house-constructed, and the largest existing, pan-cancer database of paired scRNA/TCR-seq profiles from 2.6 million T cells. Our framework is trained on data from more than 1.5 million T cells, obtained from human cancer patients across 963 tumor and blood samples, representing 14 different cancer types. Tested on additional single-cell data of more than 1 million T cells from 482 samples external to the model, we showed that scXpand provides a robust and accurate pan-cancer detection of T-cell clonal expansion from scRNA-seq data without paired TCR sequencing. It demonstrates high performance across tumor and blood samples separately, as well as across different subtypes of T cells. In addition, we created a user-friendly documentation of our framework, enabling users to directly apply it on any given dataset, as well as modifying it for other, more complex classification tasks, using our built-in optimization procedure. This setting enables a flexible foundation for broader applications according to the user preferences. Moreover, our framework provides scalable processing for handling millions of single cells with memory-efficient data streaming from disk during training, with accessibility to the already trained and optimized models for direct application of the platform.

## Results

### Constructing a comprehensive pan-cancer scRNA/TCR-seq database of T cells

To train a model for prediction of T-cell clonal expansion at the pan-cancer level, we first collected publicly available data of tumor and blood samples obtained from cancer patients across 14 cancer types, consisting of 38 datasets with more than 2.6 million T cells passing a strict quality control done separately on both scRNA-seq and scTCR-seq data^5,6,13–46^ (Figure 1A, Table S1, Method details). Overall, 26 datasets containing 963 samples with more than 1.5 million T cells were used to train the model, with additional 1 million T cells across 482 samples from 12 other datasets, used to test the model performance on each dataset independently (Figure 1A, Table S1). Across the train set, 11,950 genes shared between all datasets passed the quality control process and were used for a further integration of all datasets (Figure 1B, C, Method details). Markov Affinity-based Graph Imputation of Cells (MAGIC)^47^ was first applied in order to detect possible drop-outs of *CD8A/B*, *CD4* or *FOXP3* (Figure S1A-D, Method details). Following, single cells were labeled according to their membership in expanded or non-expanded T-cell clones, using CDR3 sequence identity, and their clone size per sample was then quantified accordingly (Figure 1B, C, Method details). This database was then prepared as an input to the model, such that the input features include a gene expression matrix containing unique molecular identifier (UMI) counts for each T cell, with gene representation using Ensembl IDs^48^ for generalization purposes and simpler user interface. The target binary variable of each T cell is its membership in either ‘expanded’ or ‘non-expanded’ T-cell clone. In addition, the tissue of origin for each T cell (i.e. tumor or blood), as well as the T-cell subtype following cell imputation (i.e. CD8^+^, CD4^+^, Treg, Double-negative or Double-positive), were also included as part of the model optimization process (Method details). Importantly, we provide an open-source, user-friendly tutorial, demonstrating how to properly prepare the model input.

**Figure 1.**
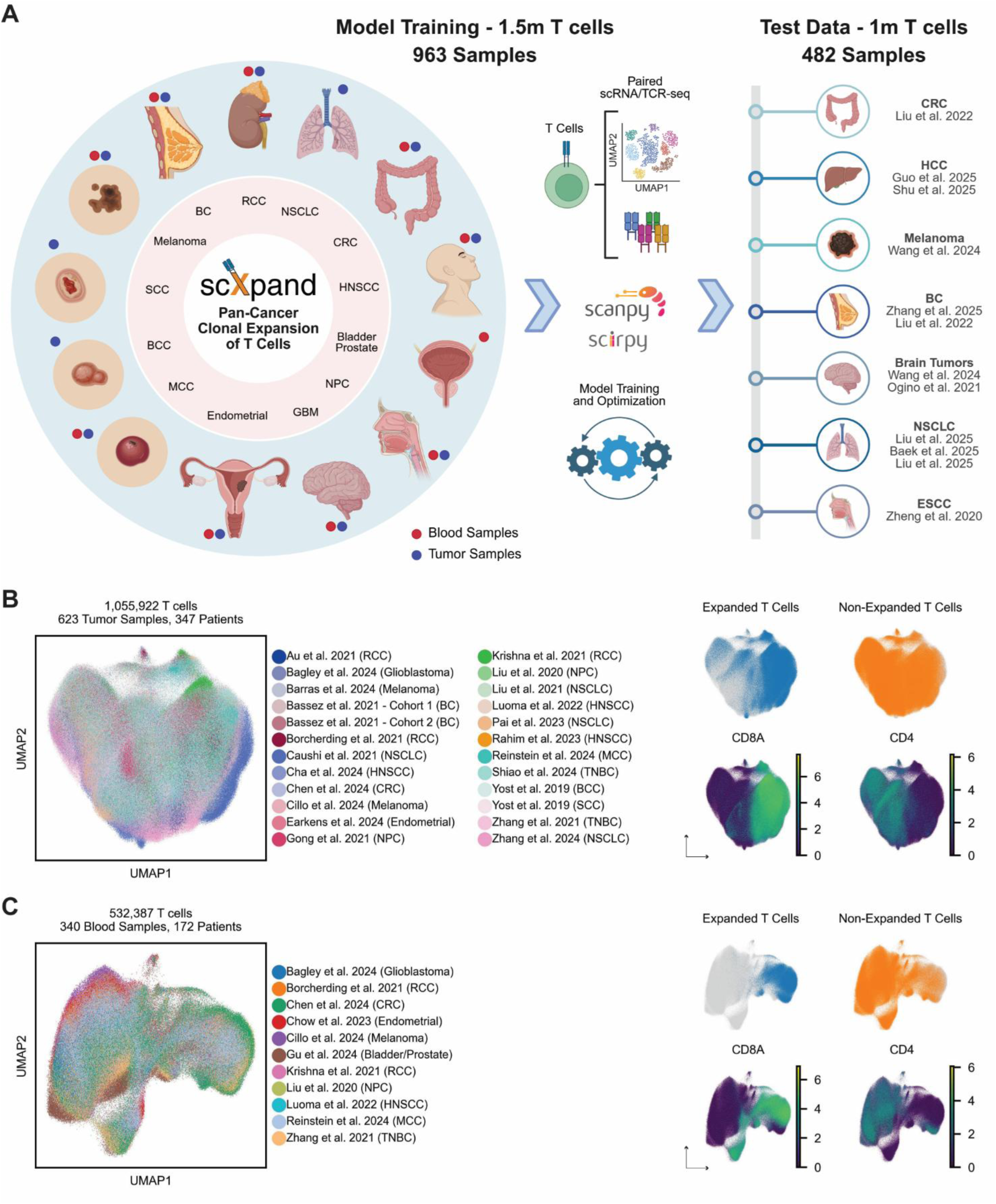
Constructing a comprehensive pan-cancer scRNA/TCR-seq database of T cells. A. A schematic workflow of the study depicting T-cell data collection across tumor and blood samples of human cancer patients, strict quality control, model training, and application on additional test datasets. B. Uniform manifold approximation and projection (UMAP) plot of 1,055,922 T cells from tumor samples having paired scRNA/TCR-seq used for model training (left), and annotations for clonal expansion as well as expression of *CD8A* and *CD4* (right). C. UMAP plot of 532,387 T cells from blood samples having paired scRNA/TCR-seq used for model training (left), and annotations for clonal expansion as well as expression of *CD8A* and *CD4* (right). See also Figure S1 and Table S1.

### Prediction of T-cell clonal expansion using scXpand

The scXpand framework for prediction of T-cell clonal expansion can accommodate different modeling strategies. In this work, we evaluated five representative models: two deep-learning models, one tree-based model, and two linear models (Method details). Alongside standard off-the-shelf methods, we developed a masked multi-task, forked count autoencoder specifically tailored to the properties of scRNA-seq data. This autoencoder denoises the input by modeling the count distribution and sparsity using a zero-inflated negative-binomial noise model, similar to Eraslan et al.^49^. On top of this autoencoder core, we added two additional branches stemming out of the latent vector to form a multi-task model. The first branch includes an auxiliary classification task of identifying the T-cell tissue of origin (i.e. tumor or blood), with the second branch consisting of an integrated classification head for the main task of predicting T-cell clonal expansion (Figure 2). This multi-task approach learns tasks in parallel while using a shared representation, potentially improves the task-specific performance^50^. As no previous models for prediction of T-cell clonal expansion exist, we compared the performance of this model with four additional models, representing widely adopted baseline models of different families: multilayer perceptron (MLP), LightGBM^51^, support vector machine (SVM), and logistic regression (Figure 3, Table S2).

**Figure 2.**
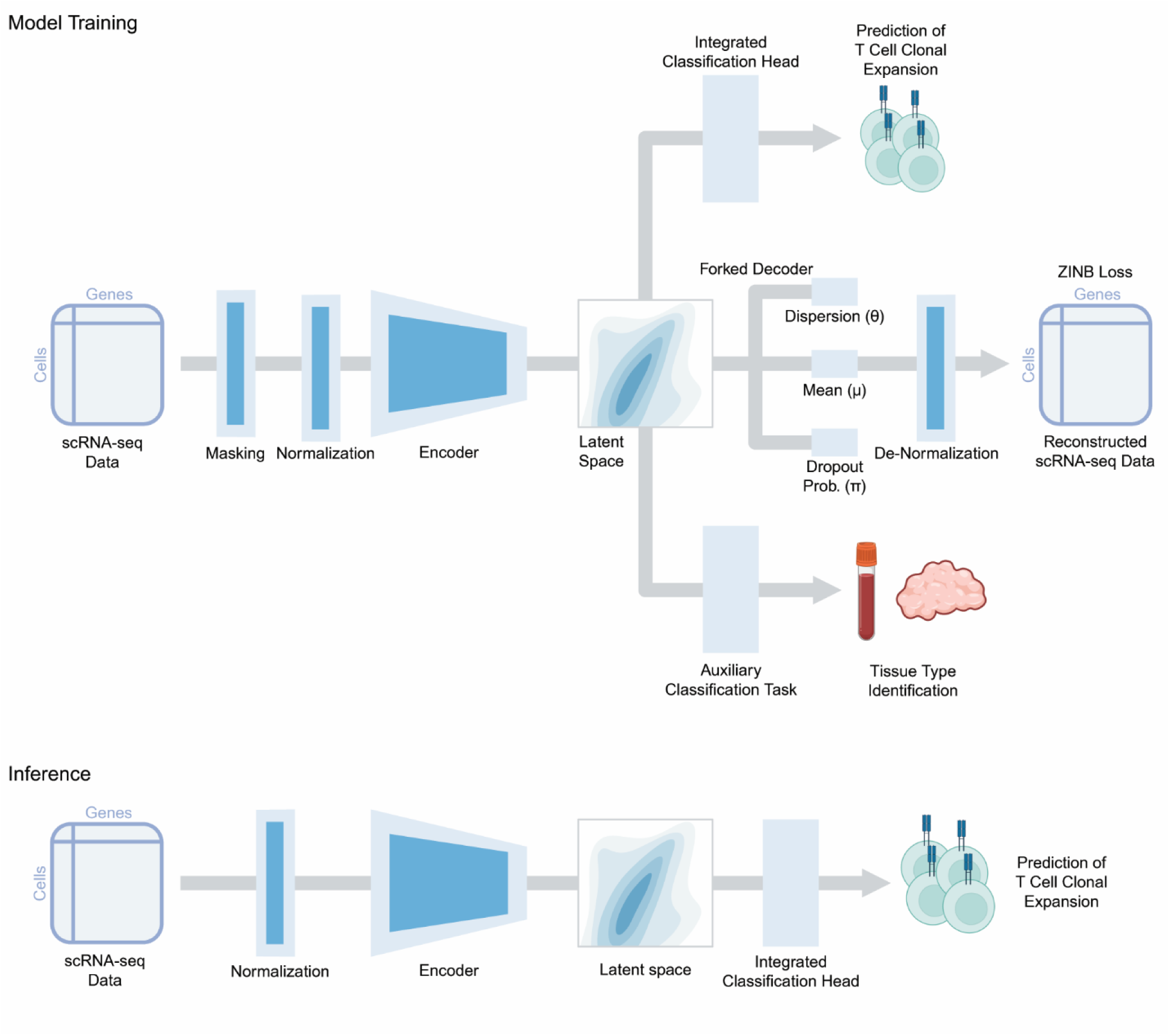
The multi-task autoencoder model architecture. A schematic description of the autoencoder architecture, demonstrating both training (top) and inference (bottom) configurations. scRNA-seq data undergo masking and normalization, with a subsequent forked autoencoder core having a zero-inflated negative binomial reconstruction loss function. Two integrated classification heads are stemming out of the latent vector, with one auxiliary classification task, and the second with the main classification task of predicting T-cell clonal expansion. Inference is being done using the main classification task alone. See also Table S2.

**Figure 3.**
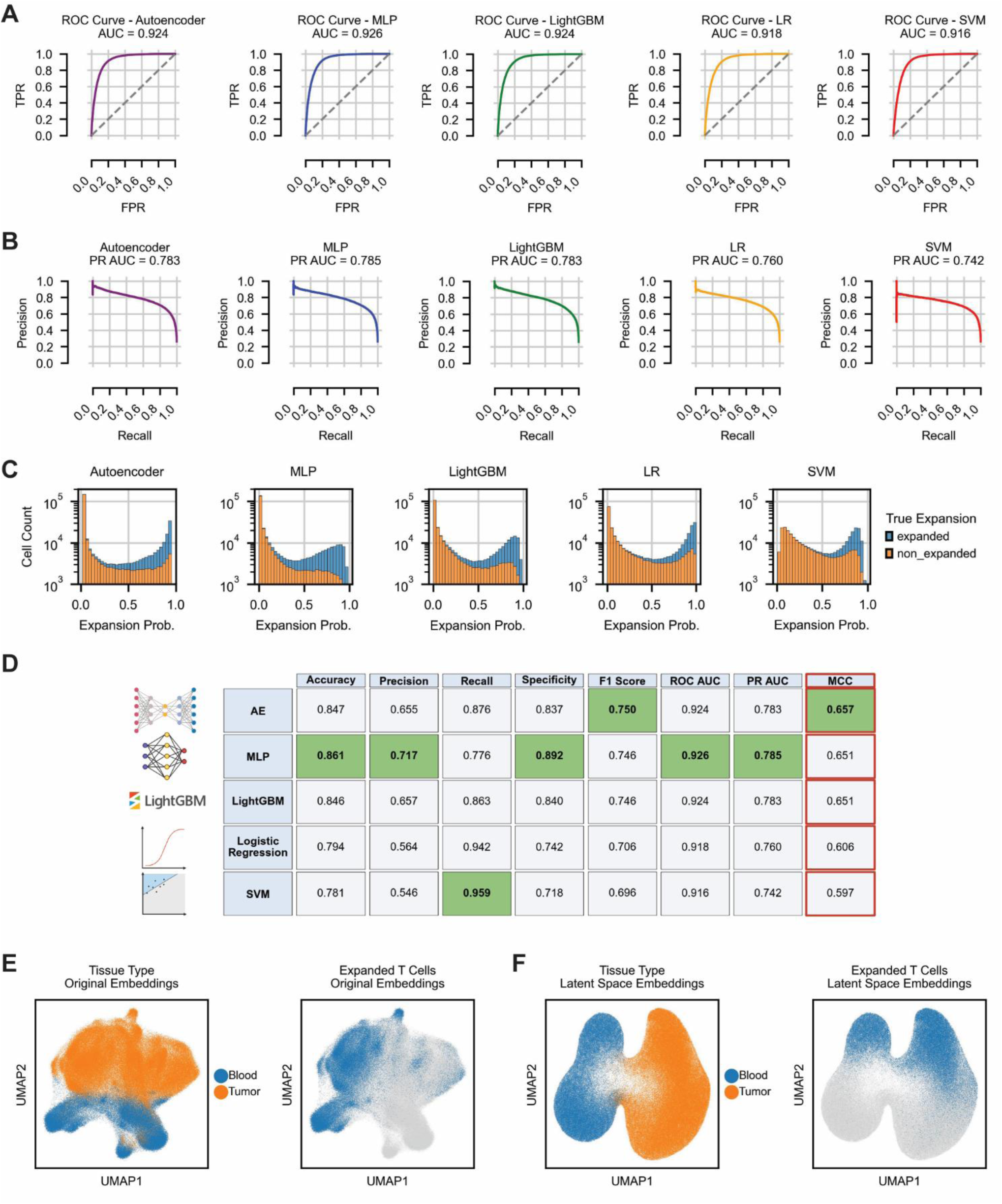
Prediction of T-cell clonal expansion using scXpand. A. Receiver operating characteristic (ROC) curves, with calculation of the corresponding area under the curves (AUCs), for each one of the five trained and optimized models, using the validation set. B. Precision-recall (PR) curves, with calculation of the corresponding area under the curves (AUCs). C. Histograms depicting the number of T cells from the validation set, across the predicted expansion probabilities obtained for each T cell by each model. Bars are colored and stacked based on the true expansion labels of each T cell. D. Model performance metrics measured using the validation set, across all five trained models. The Matthews correlation coefficient (MCC) is marked with red color. E. UMAP plot of T cells from the validation set, obtained using the common practice for dimensionality reduction and colored by tissue type (left), and T cells labeled as ‘expanded’ (right). F. UMAP plot of T cells from the validation set, obtained using the latent space embeddings of the autoencoder model, colored by tissue type (left), and T cells labeled as ‘expanded’ (right). Abbreviations: FPR, false positive rate; TPR, true positive rate; MLP, multilayer perceptron; LR, logistic regression; SVM, support vector machine. See also Figure S2 and Table S2.

From the train set of 1.5 million T cells, we allocated 80% of the data for training and 20% for validation used for hyperparameter tuning (Method details). Model performance was then evaluated on the validation set (Figure 3). Of note, we employed a stratified data allocation strategy to ensure that cell populations were proportionally represented in both the training and validation sets, while enforcing that all cells from a given patient are assigned exclusively to either training or validation set, but not both (Method details). For each optimized model we constructed the receiver operating characteristic (ROC) curve, the precision-recall (PR) curve, and calculated the area under these curves (AUC). Interestingly, the performance was similar across the different models, with no substantial differences observed in the area under the ROC (0.916-0.926, Figure 3A, Table S2) or the PR curves (0.742-0.785, Figure 3B, Table S2). Because ROCAUC and PRAUC only capture ranking performance across thresholds, they can obscure meaningful differences in classification accuracy at a fixed threshold. To overcome this limitation, we used the predicted expansion probability distribution of each model (Figure 3C) to assess accuracy, precision, recall, specificity, and F1 score, with a naive expansion probability threshold of 0.5 (Figure 3D). These additional measures revealed clear differences: some models achieved higher precision but at the cost of lower recall, whereas others showed the opposite trend. Specifically, while the autoencoder, MLP and LightGBM models showed lower false positive rates (0.109-0.163, Table S2), both the logistic regression and the SVM models demonstrated higher false positive rates (0.258-0.282, Table S2). These rates were mirrored by higher and lower false negative rates, respectively (Table S2). These results highlight that despite similar overall discrimination, the models varied in their ability to correctly identify either expanded or non-expanded T cells.

To further evaluate the model performance, and since there is a general imbalance of expanded and non-expanded T cells in the data, we calculated an additional metric, the Matthews correlation coefficient (MCC), which takes this imbalanced populations into account^52,53^ (Figure 3D). In order to address optimal expansion probability thresholds in each model, we calculated the MCC value distribution in each model per each T-cell subtype for both tumor and blood samples, separately (Figure S2A-E). This analysis resulted with different optimal thresholds per model, though MCC values remained comparable across models (Figure S2A-E). These results emphasize the robustness of the different models in capturing clonal expansion despite data imbalance and show that while the optimal thresholds may vary across models, their relative performance remains consistent.

Taken together, the scXpand platform enables robust detection of T-cell clonal expansion and lays the foundation for broader applications in immuno-oncology and cancer research. Furthermore, both the complex and the simpler models show relatively comparable performance using their optimal thresholds. This convergence underscores the robustness of the underlying biological signal, further reassuring that the observed predictions reflect genuine biological phenomena.

### Application of the model on external test datasets

As the overall MCC score for a naive threshold of 0.5 was the highest for the autoencoder model (Figure 3D), we used it as the main model for the following downstream analysis and application on the additional external 12 test datasets as described throughout the upcoming sections. Moreover, the autoencoder model provides an additional feature of data representation using its latent space embedding, which results with a smoother two-dimensional representation compared to that achieved with common dimensionality reduction methods (Figure 3E-F, Method details).

Following, we applied it on additional 12 test datasets having paired scRNA/TCR-seq, consisting of 482 samples with more than 1 million T cells (Figure 1A, Figure 4A, Table S1). Importantly, all of these datasets were external to the model and were not used as part of the training and hyperparameter tuning procedures. Applying the autoencoder model on datasets having either tumor or blood samples, as well as on datasets having both sample types, demonstrated robustly accurate results (Figures 4B-E, S3-S5, and Table S3). Specifically, on T cells obtained from tumor samples of non-small cell lung cancer (NSCLC) patients^43^, our model achieved an AUC of 0.886 (Figure 4B). Testing its performance on blood samples of melanoma patients^38^ it achieved an AUC of 0.937 (Figure 4C). In two additional datasets of tumor samples from triple-negative breast cancer (TNBC) patients^39^ and blood samples of glioma patients^42^, our model achieved an AUC of 0.877 and 0.961, respectively (Figure 4D-E). High AUCs (0.847-0.963) were obtained for additional eight datasets of brain tumors^35^, colorectal cancer^36^, lung cancer^45,46^, esophageal cancer^40^, breast cancer^41^, as well as two datasets of hepatocellular carcinoma (HCC) patients^37,44^, despite not having been trained on the latter cancer type (Figures S3-S5). As expected, and consistent with the training set, the other models exhibited comparable overall performance across datasets with a similar trade-off between precision and recall (Table S3). Overall, our five-model framework demonstrates robust and accurate performance across different datasets and cancer types, including both tumor and blood samples. These results highlight its strong generalizability and imply of its applicability in diverse contexts at the pan-cancer level.

**Figure 4.**
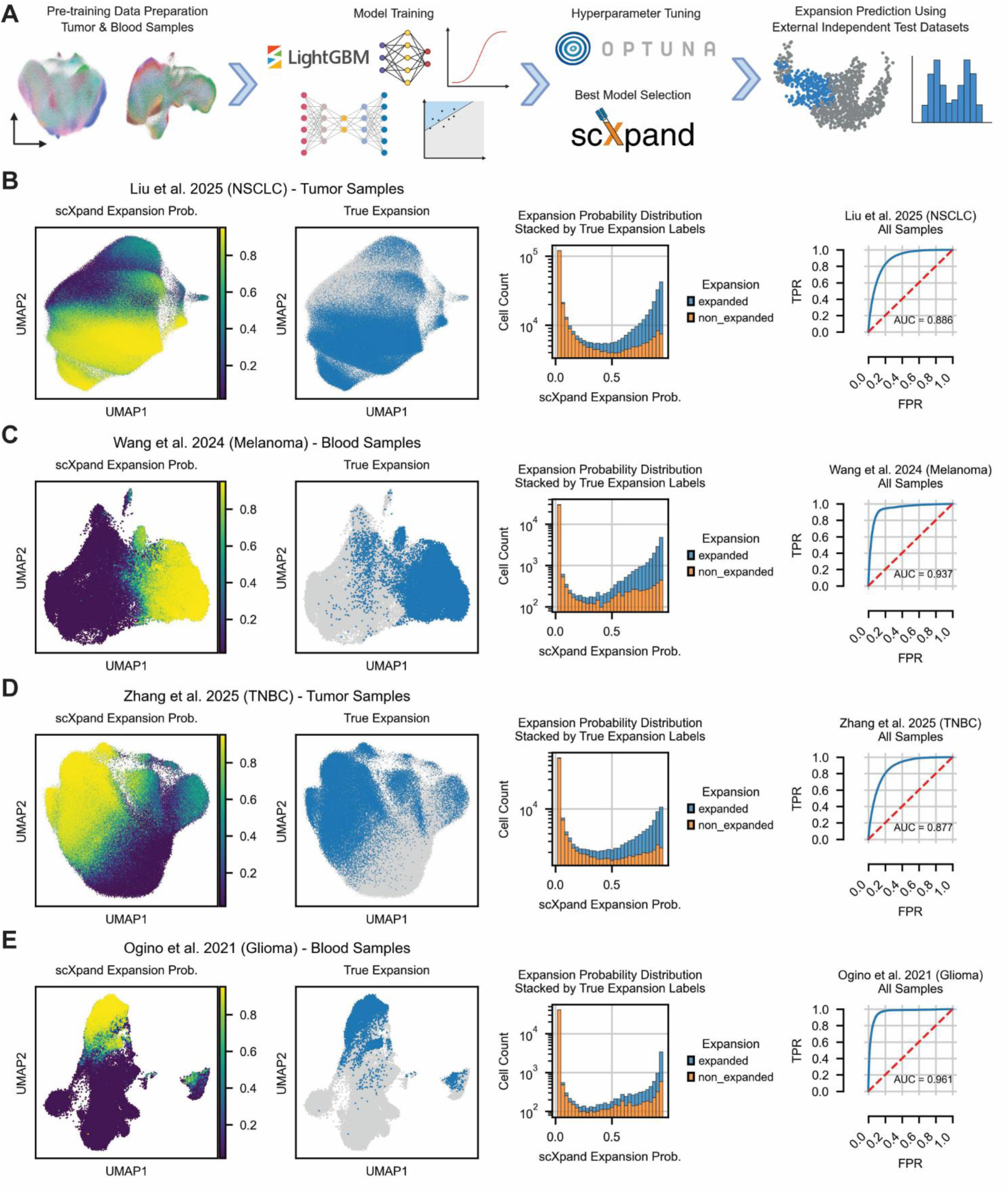
Application of the model on external test datasets. A. A schematic workflow demonstrating the model training and hyperparameter tuning processes, with application of the optimized trained models using an additional collection of test datasets that were external to the model and were not used for training. B-E. Application of the trained autoencoder model on four different test datasets^38,39,42,43^, consisting of different cancer types across tumor and blood samples. UMAP plots demonstrating the predicted expansion probability for each T cell, as well as the true expansion labels (left), including a histogram of the probability distribution stacked by the true expansion labels, with the corresponding ROC curves (right). Abbreviations: FPR, false positive rate; TPR, true positive rate. See also Figures S3-S5 as well as Tables S1 and S3.

### Model explainability and biological interpretation

In order to further understand the model output and obtain biological interpretation of its predictions, we employed the SHapley Additive exPlanations (SHAP) values^54^ that quantify the contribution of individual genes to the predicted clonal expansion status of each T cell, measuring both the magnitude and directionality of their effect on the model predictions. Furthermore, this approach reveals more intricate context-dependent gene behavior, capturing effects of gene-gene interactions and their influence on the model predictions, allowing us to identify key features highly associated with expanded or non-expanded T-cell clones.

Because SHAP value computation has been directly integrated into the LightGBM code base, allowing for exact computation of SHAP values for this method^55^, we employed SHAP value computation for the LightGBM model (Figure 5A). Among the genes having the highest absolute Shapely scores, high expression of genes such as *NKG7*, *CD8A*, *GZMA* and *CST7* was driving the model predictions towards ‘expanded’ status, while high expression of gene such as *LTB*, *SELL, CCR7, LEF1, MAL*, and *EEF1A1* was driving the predictions towards the ‘non-expanded’ status (Figure 5B). The list of genes was additionally ranked according to their global feature importance as derived from the LightGBM ‘gain’ metric, reflecting the overall contribution of each gene to the predictive performance of the model (Figure 5C).

**Figure 5.**
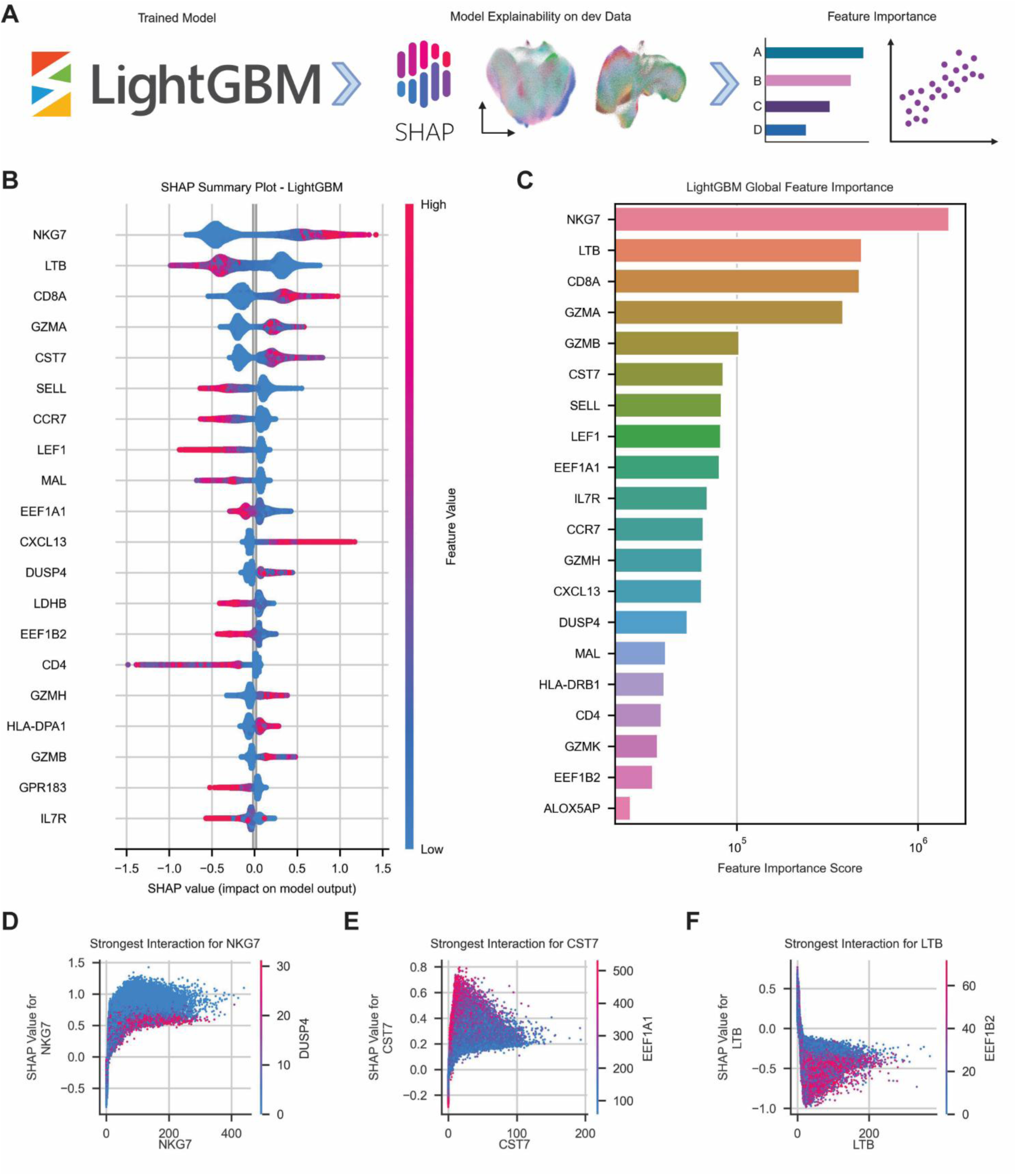
Model explainability and biological interpretation. A. A schematic workflow demonstrating the application of SHAP^54^ on the LightGBM model, using the validation set. B. SHAP summary plot depicting the top 20 important genes driving the LightGBM model predictions. C. Global feature importance of the LightGBM model, reflecting the overall contribution of the top 20 important genes to the predictive performance of the model. D-F. SHAP dependency plots showing the most interactive partners of selected genes, including *NKG7*, *CST7* and *LTB*. See also Figures S6-S7.

Indeed, in our previous work we conducted a single-cell meta-analysis of T cells in the context of cancer patients treated with checkpoint immunotherapy, and showed that expanded T cells were predominantly CD8^+^, both in tumor and blood samples^8^. In addition, differential gene expression between expanded and non-expanded T cells, demonstrated that *NKG7* was the top differentially expressed gene of expanded T cells for both tissue types^8^. Moreover, *NKG7* was previously shown to be upregulated in circulating effector/memory CD8^+^ T cells of responders to checkpoint immunotherapy compared to non-responders^56^, with induced expression of *NKG7* resulting with increased presence of intra-tumoral T cells^57^. In contrast, *LTB* was the top differentially expressed gene of non-expanded T cells for both tumor and blood samples^8^, and was previously shown to be expressed in naive-memory CD8^+^ T cells^58^. Importantly though, these genes alone cannot be used for prediction, as ∼12% of the expanded T cells do not express *NKG7*, while ∼30% of them express *LTB*. In addition, ∼23% of the non-expanded T cells do not express *LTB*, while ∼28% of them express *NKG7*, as measured for the validation set. Applying a separate SHAP value computation for CD8^+^ and CD4^+^ T cells from tumor samples, resulted with modified, yet similar list of genes driving the model predictions (Figure S6), thus showing genes driving expansion status across T-cell populations, as well as more subtype-specific genes.

Going beyond the impact of single genes and their effect on the model predictions, we next examined SHAP values for gene-gene interactions. To this end we focused on the top genes having the highest absolute Shapely scores, and examined their strongest interactive partners (Figure 5D-F). This process resulted with notable gene pairs demonstrating context-dependent effect, such that the SHAP value for a gene can be different between two cells having the same expression level of that gene, resulting from the contextual expression of other genes in these cells. More specifically, while high expression of both *NKG7* and *DUSP4* resulted with higher SHAP values separately (Figure 5B), the interaction between both genes demonstrates that high expression of *DUSP4* in the context of *NKG7* expression, reduces SHAP values (Figure 5D). In contrast, while high expression of *EEF1A1* resulted with low SHAP values (Figure 5B), high expression of *EEF1A1* in the context of *CST7* expression can result with increased SHAP values (Figure 5E). Moreover, while high expression of both *LTB* and *EEF1B2* resulted with lower SHAP values (Figure 5B), their contextual expression demonstrates synergistic effect in lowering the SHAP values (Figure 5F). As expected, SHAP value computation using a simple model such as the logistic regression model, resulted with capturing of simple linear trends between gene expression and SHAP values (Figure S7A-B). These results emphasize the tradeoff between model complexity, interpretability, and the ability to capture more complex context-dependent interactions between features. Overall, by using SHAP for model explainability we demonstrate the capturing of previously known biological effects of gene expression and T-cell clonality, as well as more intricate, non-trivial gene-gene interactions affecting model predictions.

### Applicable examples of using the scXpand framework

In order to further demonstrate the applicability of our platform, we applied the autoencoder model on a publicly available dataset of breast cancer patients having scRNA-seq data alone, without paired scTCR-seq data^59^ (Figure 6A, S8A, Table S1, Method details). This dataset consists of 14 patients, with gene expression data of 9 cell types, including T cells (Figure 6A). In addition, patients were originally divided by the authors according to two types of immune environments (IEs), based on the cellular state of T cells: 1. an exhausted environment, IE1, with presence of PD-1^high^/CTLA-4^high^/CD38^high^ T cells, and 2. Non-exhausted environment, IE2, containing T cells that did not express exhaustion markers^59^ (Figure 6A). By applying our model on T cells alone, we identified those having high probability of being expanded (Figure S8A), and labeled the corresponding CD8^+^ T cells as ‘expanded’ (Figure 6A, Method details). We then quantified the abundance of expanded CD8^+^ T cells, out of all the CD8^+^ T cells per sample (Figure 6B), and compared this abundance between both immune environments. Although the overall T-cell frequency was previously shown to be similar in both groups^59^, we found that expanded CD8^+^ T cells are significantly more abundant in IE1 compared to IE2 (p-value = 0.025, Figure 6C), suggesting of the potential connection between the immune environment and T-cell clonal expansion. We then used our previously devised 12 gene signature of T-cell clonal expansion in the context of patient response to checkpoint immunotherapy^8^, and applied it on expanded CD8^+^ T cells per sample (Method details). This signature was previously used by us to link clonal expansion to either improved or poor clinical outcomes across multiple cancer types^8^. Applying this signature on the breast cancer dataset resulted with a score per sample between 0 and 1, quantifying the amount of expanded CD8^+^ T cells associated with good prognosis, out of all the expanded CD8^+^ T cells per sample (Figure 6D). Comparing this score between both immune environments showed significant difference between IE1 and IE2 (p-value = 0.009, Figure 6D), showcasing that the cellular state of expanded CD8^+^ T cells is also different between both environments, in addition to the difference in the general abundance of expanded CD8^+^ T cells. Overall, this analysis demonstrates the applicability of our model on a single-cell dataset lacking TCR sequencing, and its ability to provide additional valuable information for the downstream data analysis.

**Figure 6.**
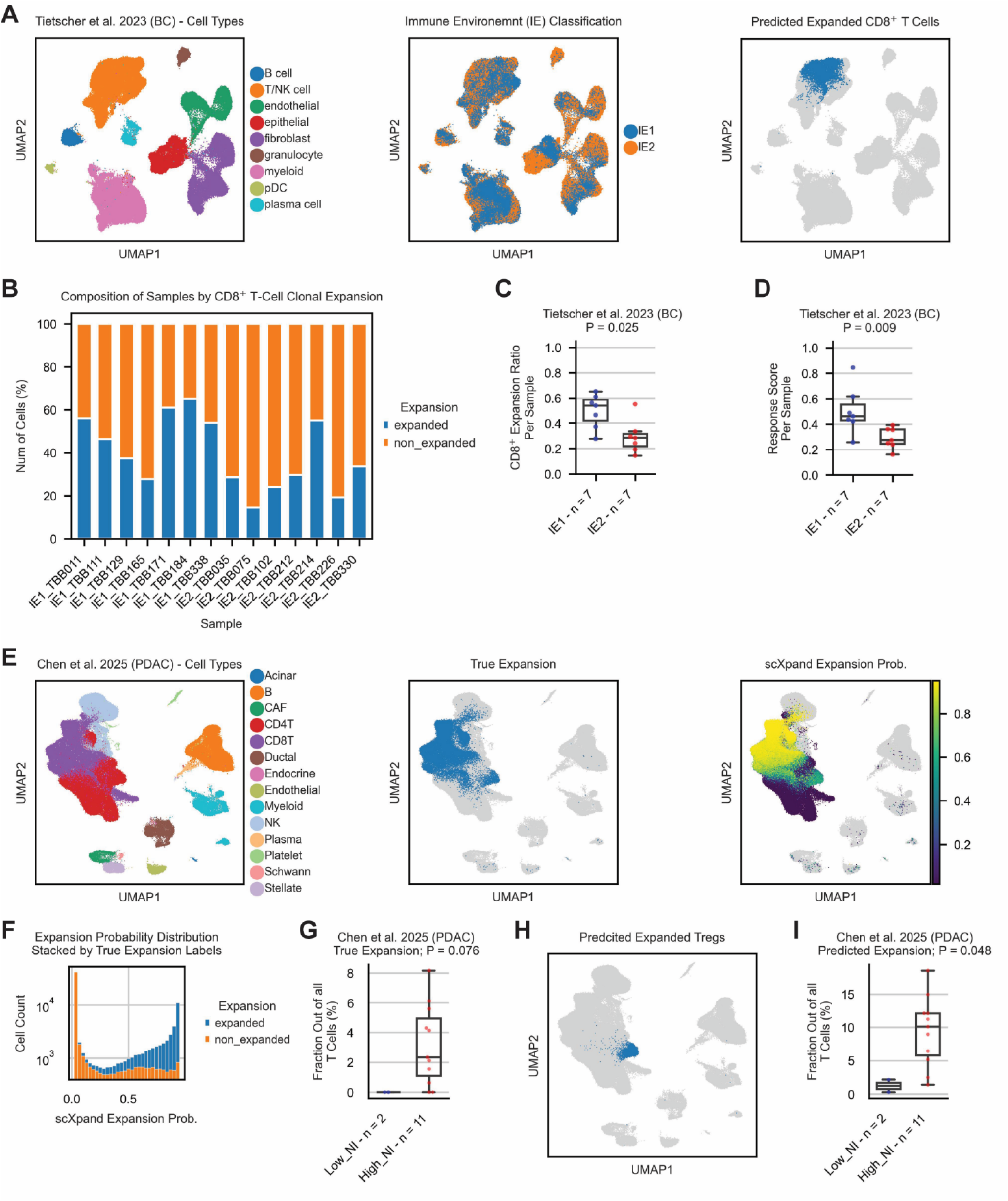
Applicable examples of using the scXpand framework. A. UMAP plots of single cells from a publicly available dataset of breast cancer patients^59^, colored by attribution of single cells to cell types and immune environment (IE) classification, as originally provided by the authors^59^, as well as labeling of CD8^+^ T cells predicted as expanded by the autoencoder model. B. Stacked barplot showing the fraction of predicted expanded CD8^+^ T cells out of all the CD8^+^ T cells per sample. C. Boxplot showing the comparison of expanded CD8^+^ T-cell fraction per sample, between the different immune environments (p-value = 0.025, n=14). D. Comparison of a response signature devised by Shorer et al.^8^, using expanded CD8^+^ T cells, between the different immune environments (p-value = 0.009, n=14). E. UMAP plots of single cells from a publicly available dataset of PDAC patients^60^, colored by attribution of single cells to cell types as originally provided by the authors^60^, as well as labeling of truly expanded T cells based on scTCR-seq data and the predicted expansion probability of the autoencoder model. F. A histogram of the predicted expansion probability distribution, stacked by true expansion labels. G. Boxplot showing the comparison of truly expanded Treg cell fraction based on scTCR-seq data, out of all the T cells per sample, between tumor samples having different cancer-neural invasion status (p-value = 0.076, n = 13). H. UMAP plot of single cells colored by the labeling of intra-tumoral Treg cells predicted as expanded by the autoencoder model. I. Boxplot showing the comparison of predicted expanded Treg cell fraction by the autoencoder model, out of all the T cells per sample, between tumor samples having different cancer-neural invasion status (p-value = 0.048, n = 13). Abbreviations: BC, breast cancer; PDAC, pancreatic ductal adenocarcinoma; NI, cancer-neural invasion; p-values were calculated using a two-sided Wilcoxon rank-sum test. See also Figure S8 and Table S1.

To further demonstrate the generalizability of our framework, we applied the autoencoder model on an additional publicly available dataset of pancreatic ductal adenocarcinoma (PDAC) patients, having paired scRNA/TCR-seq^60^ (Figure 6E, S8B, Table S1, Method details). This dataset includes 29 samples of PDAC patients passing our quality-control, labeled according to their cancer-neural invasion (NI) status as provided by the authors^60^, including adjacent normal, tumor, and blood samples (Figure S8C). Using its available scTCR-seq data, we measured the performance of the model in predicting T-cell clonal expansion (Figure 6E-F), and calculated the area under the ROC curves for each sample type (Figure S8D). Our model achieved an area under the ROC curve of 0.868, 0.969, and 0.935, across tumor, blood, and adjacent normal samples, respectively (Figure S8D). These results emphasize the generalizability of our framework, as both pancreatic cancer and adjacent normal samples were not considered during the training process. Interestingly, regulatory T cells (Tregs) were observed by the authors to be in higher proportions within tumor samples of patients having high NI status^60^. Using the available scTCR-seq data, we managed to show that this finding corresponds to a higher proportion of expanded Tregs out of all the T cells per sample in the high NI group (p-value = 0.076, n = 13, Figure 6G). Importantly, by predicting expanded Tregs in tumor samples using our model (Figure 6H), we managed to similarly regenerate this finding (p-value = 0.048, n = 13, Figure 6I). Taken together, these results demonstrate the generalizability of our model across multiple tissue types and subtypes of T cells at the pan-cancer level, with the ability to be applied on datasets having scRNA-seq data alone, without paired scTCR-seq.

## Discussion

The anti-tumor activity of T-cell clones has been widely studied in the context of cancer patient response to therapy, using paired scRNA/TCR-seq data. However, up to date, there is currently no existing model for prediction of T-cell clonal expansion from scRNA-seq data without paired scTCR-seq data. Our scXpand platform is bridging this gap by providing a set of optimized models that were trained and validated using our in-house-constructed, and the largest existing comprehensive human pan-cancer database of T cells, across both tumor and blood samples. Our platform provides a user-friendly, well documented interface that can be easily used for application, training, and optimization of machine learning models, enabling to handle millions of single cells in a memory-efficient manner. Validating the performance of our platform on a set of 12 test datasets external to the model, we showed the robustness, accuracy and generalizability of our results across different datasets and cancer types. In addition, we showed the main important genes driving the model decisions and depicted non-trivial gene-gene interactions affecting the model predictions. We also provided biological interpretation of these results and showed the applicability of our platform on a breast cancer dataset having scRNA-seq data alone, as well as on a PDAC dataset having paired scRNA/TCR-seq data.

As part of the scXpand platform we also introduced a multi-task masked autoencoder which takes advantage of the specific and unique properties of scRNA-seq data, such as its sparsity and count distribution. We used this model to obtain a new 2D representation of the data using its latent space embedding, and used its integrated classification head for detection of T-cell clonal expansion. Importantly, all of the trained optimized models are publicly available and are ready for direct application or de-novo training by the user, including possible user-made modifications to handle other single-cell classification tasks.

Although our platform was trained on data at the pan-cancer level, the training process was done on 14 cancer types while excluding other important cancer types, such as pancreatic, ovarian, gastric, and cervical cancer, as well as hematologic malignancies. Indeed, more cancer types can be addressed in future studies as more relevant scRNA/TCR-seq data become available. However, we did show that our platform performs well even on cancer types that were not included during the training process, such as hepatocellular carcinoma (HCC)^37,44^ and pancreatic cancer^60^. In addition, the model was trained using only tumor and blood samples, without other tissue types such as lymph node samples and adjacent normal tissue. Yet, applying our model on the PDAC data by Chen et al.^60^, we showed that high performance can be achieved on adjacent normal samples as well. Importantly, and corresponding with previous studies analyzing different single-cell models in various tasks^61,62^, all five models trained within our platform showed relatively comparable overall performance despite varying levels of complexity. However, they differed in their ability to correctly identify either expanded or non-expanded T cells, emphasizing their inherent trade-offs and highlighting the importance of selecting models based on the specific biological question or application. Moreover, the more complex models, such as LightGBM, capture the more intricate gene-gene interactions, providing non-linear trends between gene expression and model predictions. Finally, it is important to note that our platform does not address the actual clonal identity of T cells nor their TCR sequence identity, thus limited only to detection of a cellular state of T-cell clonal expansion based on gene expression. Nonetheless, our platform successfully links scRNA-seq data to clonal expansion, providing the first existing framework for this task. This serves as a foundation for future frameworks and additional scalable single-cell classification tasks in favor of the scientific community worldwide, with a special dedication to the cancer research community.

## Resource availability

### Lead contact

Further information and requests for resources should be directed to and will be fulfilled by the lead contact, Keren Yizhak.

### Materials availability

This study did not generate new unique reagents.

### Data and code availability

- This paper analyzes existing, publicly available scRNA/TCR-seq data. The accession numbers for these datasets are summarized in Table S1.
- All original code has been deposited at https://github.com/yizhak-lab-ccg/scXpand and is publicly available as of the date of publication. The complete set of trained models is publicly accessible via Figshare^63^ (https://doi.org/10.6084/m9.figshare.30067666). The full documentation of our platform, including tutorials and guidelines, is available on Read the Docs (https://scxpand.readthedocs.io). An official python package is provided via PyPi (https://pypi.org/project/scxpand), with an additional GPU-supported version (https://pypi.org/project/scxpand-cuda).
- Any additional information required to reanalyze the data reported in this paper is available from the lead contact upon request.

## Acknowledgments

This work was supported by the Israel Science Foundation (1124/25), the Israel Cancer Research Fund (23-204-RCDA) and the Eric and Wendy Schmidt Fund for Strategic Innovation. This work received additional support by the Ruth and Bruce Rappaport Technion Integrated Cancer Center (RTICC). Figures 1A, 2, 3D, 4A, and 5A, were created with BioRender.com using a paid license.

## Author Contributions

O.S, R.A and K.Y conceived the idea. O.S, R.A and K.Y designed the study. O.S and R.A performed the analysis. O.S, R.A and K.Y wrote the manuscript.

## Declaration of interests

The authors declare no competing interests.

## Experimental model and study participant details

This study did not involve any new experimental models, human participants, or animal subjects. All data used in this study are publicly available, with samples and single cells obtained from each dataset described in Table S1 following our quality control. Details regarding data acquisition and processing are provided in the method details.

## Method details

### Acquisition of paired scRNA/TCR-seq datasets and preprocessing

26 single-cell datasets of cancer patients having paired scRNA/TCR-seq data were collected together with their annotations for clinical outcomes and treatment time-point, where available. 15 datasets contained tumor biopsies, 9 datasets contained both tumor and blood samples, and 2 datasets contained blood samples alone (Table S1).

All scRNA-seq datasets were droplet-based and contained unique molecular identifier (UMI) counts. For each dataset, we followed similar preprocessing steps to those we previously conducted^8^ using Scanpy^64^: We removed cells expressing less than 200 genes and removed genes that were expressed in less than 3 cells. We also removed cells having more than 10% of mitochondrial gene-count. In the dataset of Krishna et al.^15^, scRNA-seq was provided following the authors’ original quality-control (QC) while excluding mitochondrial genes from the provided gene expression matrix. In that case only, we used the expression matrix as provided by the authors following their QC, retaining cells with less than 20% of mitochondrial gene-count. We also applied Scrublet^65^ to remove potential doublets and filtered-out cells having *‘doublet_score’* larger than 0.3 and/or those predicted as doublets with score below 0.3.

For the scTCR-seq datasets, we rearranged each dataset to be compatible with the Adaptive Immune Receptor Repertoire (AIRR) schema that was further preprocessed by Scirpy^66^ and could be directly uploaded using the *‘scirpy.io.read_10x_vdj’* function. For each scTCR-seq dataset, we applied quality control based on the standard protocol suggested by the authors. In short, we removed cells having multi-chains, orphan VJ or orphan VDJ chains.

### Integration of scRNA/TCR-seq datasets

Following the quality control process described above, we removed samples having less than 3 cells overall, and integrated all processed datasets resulting with a total of 1,055,922 single cells from tumor samples and 532,387 single cells from blood samples, all with paired scRNA/TCR-seq. Overall, 11,950 genes existed across all datasets and were used for further analysis, such that tumor and blood samples were visualized separately.

For visualization purposes only (Figures 1B-C, S1A-D), we normalized the expression level in the integrated scRNA-seq datasets to a standard target sum of 10,000 counts per cell and then applied log2-transformation. Highly variable genes were calculated using *‘scanpy.pp.highly_variable_genes’* with a *‘batch_key’* of *‘cancer type’* in order to preserve biological differences between the different cancer types. We then calculated a 40-component PCA and applied Batch Balanced K-Nearest Neighbors (BBKNN)^67^ with a ‘*batch_key’* of *‘sample’* in order to remove batch effects between samples. UMAP^68^ was used for dimensionality reduction and data visualization.

### Markov Affinity-based Graph Imputation of Cells (MAGIC) for detection of drop-outs and labeling of expanded clones

In order to identify possible drop-outs in our integrated datasets, we applied MAGIC^47^ on the log2-normalized count matrices of tumor and blood samples separately, using all the genes and single cells passing our QC, while using the default arguments of the algorithm implementation in Scanpy^64^. We first fitted a density curve to the log2-transformed imputed gene expression of *CD8A/B*, *CD4*, and *FOXP3*, and set the expression threshold for each gene as the trough of the bimodal density curve (Figure S1A-B). For the imputed gene expression of *FOXP3*, the threshold was set as 1. For the non-imputed expression of these genes, log2-transformed expression threshold was also set as 1. Overall, all the T cells that passed our QC process were further divided into five subtypes (Figure S1C-D): we considered a single cell to be CD8^+^, when the imputed or non-imputed gene expression of either *CD8A* or *CD8B* was above the threshold. Similarly, a single cell was considered to be CD4^+^, when the imputed or non-imputed gene expression of *CD4* was above the threshold. We also defined double-positive cells as those that were considered to be both CD8^+^ and CD4^+^. Double-negative cells were those that were both CD8^-^ and CD4^-^. Treg cells were defined as those that are CD4^+^ having imputed or non-imputed gene expression of *FOXP3* above the threshold. In cases where the imputed expression resulted with loss of signal from the non-imputed gene expression, we remained consistent with the non-imputed gene expression.

We then defined clonotypes based on the identity of the CDR3 nucleic acid sequence of each cell using *‘scirpy.tl.define_clonotypes’*, such that both the VJ and VDJ CDR3 sequences had to match and the T-cell subtype of all the cells in each clone is the same (i.e. CD8^+^, CD4^+^, double-positive, double-negative or Treg). In cases where more than one pair of VJ and VDJ sequences was detected per cell, we considered the most abundant pair of VJ/VDJ chains where applicable. In the dataset of Krishna et al.^15^, scTCR-seq was provided only with the amino acid sequence of the CDR3 region. In that case only, clonotypes were defined based on the amino acid identity of the CDR3 sequence.

We defined a clonotype as ‘expanded’ per sample according to Shiao et al.^18^, and as we did previously^8^, such that ‘expanded clones’ were those that contained more than 1.5x the median number of cells found in each clonotype in that sample, while excluding singletons. In order to conduct further analysis at the level of each sample, we assigned each clone with a unique ID such that the same clone appearing in multiple samples from the same patient received a different ID, indicating the clone sample of origin. We then used VDJdb^69^ as a reference database for annotating epitopes based on amino acid sequence identity according to the standard protocol suggested by the authors^66^.

### Creating the scXpand framework

We created a framework consisting of five different models. These include two linear models (SVM and logistic regression), one tree-based model (LightGBM), and two deep-learning models that include a multilayer perceptron (MLP) and a multi-task autoencoder. Both linear models of SVM and logistic regression were trained using Scikit-learn implementation of stochastic gradient descent classifier (‘*sklearn.linear_model.SGDClassifier’*)^70,71^, enabling large-scale learning with efficiency and ease of implementation. The tree-based LightGBM gradient boosting model^51^, supports fast training and high efficiency, with relatively low memory usage, enabling it to handle large-scale data. It was trained using its built-in implementation in Python (‘*lightgbm.LGBMClassifier’*). As for the two deep-learning models, we used the PyTorch library^72^ in order to define the architecture of each model. In short, the MLP model includes fully connected layers, incorporating dropout regularization to mitigate overfitting, with an optimization handled using the AdamW optimizer^73^, while a scheduler selected from multiple possible options dynamically adjusts the learning rate (Table S2). The autoencoder-based model is an extension of the deep count autoencoder (DCA) model^49^ to the supervised learning setting. The DCA model is composed of a decoder and encoder and is trained with only gene counts in an unsupervised manner, in order to learn a meaningful lower dimensional embedding of the data. The decoder results with the parameters of the zero inflated negative binomial model (ZINB): mean, dispersion, and dropout probability parameter. We expanded this model by adding another classification head module that predicts the expansion state based on the latent vector (Figure 2). All of the modules are jointly trained end-to-end with a loss function that adds the classification loss and reconstruction loss (based on the ZINB likelihood function). For the autoencoder-based model, we implemented two possible configurations of standard and forked architecture. The forked autoencoder extends the standard design by using separate decoder branches for the mean, dispersion, and dropout probability parameters, enabling more flexible modeling of complex data distributions. In addition to the integrated classification head for the main task of predicting T-cell clonal expansion, this model supports additional possible auxiliary classification tasks, such as identification of the tissue type of origin for each T cell (i.e. tumor or blood), identification of T-cell subtype (i.e. CD8^+^, CD4^+^, Treg, Double Negative or Double Positive), or both. Importantly, this model enables multiple different reconstruction loss functions, including negative binomial, zero-inflated negative binomial, or MSE.

The final architecture of each model, including the number of hidden layers, layer dimensions, the existence and structure of the auxiliary classification heads, as well as the hyperparameter tuning for all of the five different models, were determined following an optimization process using Optuna^74^, with 100 trials for each model. The full lists of hyperparameters for each model appears in Table S2, with the readily trained and optimized models that are publicly accessible.

### Model training

Each one of the different five models was trained on a set of more than 1.5 million T cells, following a structured workflow, beginning with data preprocessing and a stratified splitting into training (80%) and validation (20%) sets. Preprocessing included count normalization to a target sum of 10,000 counts per cell, including an optional log transformation (if specified), and followed by Z-score normalization, where the empirical mean and standard deviation per gene were calculated using only the training set. Data stratification ensured similar distribution of tissue types (blood or tumor) and cancer types between the train and validation sets. We also restricted all of the samples obtained from a single patient to be used in either train or validation, but not both, ensuring that data from the same patient cannot appear in both sets. During the training, we used the following two data augmentation methods: to improve robustness to sequencing drop-outs, we added random masking to the gene expression matrix, randomly zeroing small subset of the gene counts and added random white Gaussian noise to increase robustness for measurement errors. Hyperparameter tuning for each model was conducted using Optuna^74^, which explores parameter spaces across multiple trials, enabling efficient trial pruning during model optimization. The pipeline integrates automated logging, checkpointing, and systematic model evaluation to ensure reproducibility. Model inference is executed with batched evaluation on validation data, and performance is assessed using comprehensive metrics evaluated for different pairs of tissue-type (blood or tumor) and T-cell subtypes (CD8^+^, CD4^+^, double negative, double positive, and Tregs). Optimal configuration of hyperparameters was selected for each model based on the harmonic average of the AUCs across all possible ‘tissue type – T-cell subtype’ combinations. This was done in order to ensure the model performs well across all possible T-cell subtypes from the different tissue types, as well as giving higher weights to low AUC values using the harmonic average, compared to the standard arithmetic mean. Once trained, the model is saved for further downstream analyses, ensuring its applicability on external scRNA-seq datasets. Importantly, the model is compatible with Scanpy standard data structure^64^ and was trained on soft labels for T-cell expansion in the range of 0 to 1, defined by the ratio between the clone size and 1.5x the median number of cells found in each clonotype per sample. Hardware used for the training of both deep-learning models and the LightGBM model, included a workstation equipped with an Intel core Ultra 9 285K CPU 256 GB RAM and an NVIDIA RTX 6000 Ada Generation GPU 48 GB VRAM. Both linear models (logistic regression and SVM) were trained using a working station equipped with an Intel core i9-14900KF CPU 128 GB RAM and an NVIDIA GeForce RTX 4080 SUPER GPU 16 GB VRAM.

### Comparison between the different trained models

Across all 100 trials per model, we selected the best trial based on the harmonic average of the AUCs across all possible ‘tissue type – T-cell subtype’ pairs, as described above. We then compared the performance of each model on the validation set using a set of evaluation metrics, such that a T-cell was predicted as ‘expanded’ using a naive threshold of 0.5 for the expansion probability predicted by the model. These metrics include accuracy, precision, recall, specificity, F1 score, area under the ROC curve, area under the precision-recall curve, as well as the Matthews correlation coefficient (MCC)^52,53^, which considers the imbalance of labels in our data (Figure 3, Table S2). The MCC distribution by the predicted expansion probability was then calculated for each model using all possible ‘tissue type – T-cell subtype’ combinations (Figure S2), including identification of optimal probability threshold for each combination in each model.

In addition, and specifically for the autoencoder model, we also used its latent vector to obtain a new 2D representation of the data (Figure 3F). We used the 32-dimensional latent vector of the autoencoder and extracted the most highly variable dimensions with the function ‘*scanpy.pp.highly_variable_genes*’. We then applied PCA and used Batch Balanced K-Nearest Neighbors (BBKNN)^67^ with a ‘*batch_key’* of *‘sample’* in order to remove batch effects between samples. UMAP^68^ was applied on the PCA results. We compared this visualization with our regular visualization pipeline using gene expression as described above (Figure 3E). We applied this pipeline on the validation set separately for comparison purposes. A tutorial demonstrating how to obtain the latent space embedding and its visualization is provided within the platform’s documentation.

### scXpand implementation on test datasets

In order to test the performance of each model in an unbiased manner, we used additional 12 datasets containing more than 1 million T cells from 482 samples. All of these datasets were entirely external to the model and were not used during the training and optimization procedures. For each dataset, we applied a separate quality control on each of the two data modalities, scRNA-seq and scTCR-seq, as described above, and labeled the expansion status of each T cell accordingly. We then applied each model using the raw UMI count matrix of each dataset, obtaining an expansion probability for every single T cell (Figures 4, S3-S5, Table S3). In cases where genes that were used during training were missing in the test dataset (Table S3), they were automatically added to the relevant dataset with a gene expression of zero, mimicking potential sequencing dropouts, and were passed with the formatted gene expression matrix as an input to the model. For each model we measured the area under the ROC curve across all datasets and visualized the distribution of the expansion probability, stacked by the true expansion status. We used UMAP for visualization purposes, as described above.

### Model explainability using SHAP

In order to provide a biological interpretation of the model predictions, we used the SHapley Additive exPlanations (SHAP) values^54,55^ for the validation set. We used the implementation of the SHAP Python library with default parameters, and applied it on the LightGBM model. We used the ‘*shap.summary_plot*’ function in order to visualize the 20 most important features driving the model predictions, together with their directionality and magnitude of their impact (Figure 5B). We then used the ‘*shap.dependence_plot*’ function to show the effect of single genes across the entire validation set, accounting for possible interaction effects with other genes (Figure 5D-F). We did this process using all the cells in the validation set, as well as on CD8^+^ and CD4^+^ T cells from tumor samples separately (Figure S6A-B). It is important to note that the same exact LightGBM model was used for SHAP value computation for the validation set, as well as for CD4^+^ and CD8^+^ T cells from tumor samples. We repeated this process for the logistic regression model using all the cells from the validation set to demonstrate differences of gene-gene interactions between the simple logistic regression model and the more complex tree-based LightGBM model (Figure S7A-B). Specifically for the LightGBM model, we also used its built-in ‘*feature_importance*’ function with ‘*importance_type = gain*’ to reflect the overall contribution of each gene to the predictive performance of the model (Figure 5C).

### Application of the autoencoder on scRNA-seq dataset of breast cancer patients without paired scTCR-seq

We utilized a publicly available scRNA-seq dataset of 14 breast cancer patients, including cell-type classification and patient stratification by immune environments, as originally provided by the authors^59^ (Figure 6A, S8A, and Table S1). We applied a strict quality control using the gene expression matrix as described above, including filtration of cells, genes, doublet removal, followed by extraction of highly variable genes and visualization using UMAP. MAGIC^47^ was applied to detect sequencing dropouts as described above, followed by classification of T-cell subtypes. We applied our autoencoder model using only cells classified as T cells by the authors, and detected expanded CD8^+^ T cells using the relevant optimal autoencoder threshold for CD8^+^ T cells in tumor samples (Figure S2A). We then calculated the fraction of expanded CD8^+^ T cells out of all the CD8^+^ T cells per sample and used a two-sided Wilcoxon rank-sum test to compare this ratio between patients based on their immune environment classification (Figure 6B-C). Focusing on expanded CD8^+^ T cells alone, we applied our previously devised response score of expanded CD8^+^ T cells in the context of patient response to checkpoint immunotherapy^8^, and calculated a response score per patient. In short, we used a twelve gene signature, including *CXCL13*, *DUSP4*, *RBPJ*, *LYST*, *GZMK* and *HLA-DQA1*, accounting for good prognosis, as well as *FOS*, *ZNF683*, *CTSC*, *GZMH*, *ANXA1* and *XCL2*, accounting for poor clinical outcomes. We then used these two lists of genes and scored each expanded CD8^+^ T cell based on the number of expressed genes (log_2_(*normalized*_*counts* + 1) > 1) out of the 6 markers in each list, yielding two scores for each expanded CD8^+^ T cell. Then, each CD8^+^ T cell was classified as ‘favorable’ or ‘unfavorable’ based on the majority vote of the two scores. In cases where both scores were tied in a single cell, it was classified as ‘favorable’. Finally, we computed a score per patient by taking the ratio between the number of cells classified as ‘favorable’ out of the total number of cells classified as ‘favorable’ or ‘unfavorable’, yielding a score per patient between 0 and 1. CD8^+^ T cells demonstrating an expression of zero markers from both lists of markers were excluded from this analysis. We then used a two-sided Wilcoxon rank-sum test to compare these scores between patients based on their immune environment classification (Figure 6D).

### Application of the autoencoder on scRNA/TCR-seq dataset of PDAC patients

We utilized a publicly available scRNA/TCR-seq dataset of PDAC patients, including cell-type classification and patient stratification by cancer-neural invasion (NI) status, as originally provided by the authors^60^ (Figure 6E, S8B, and Table S1). We applied a strict quality control using both the gene expression matrix and the scTCR-seq data as described above, including labeling of expanded T cells with a specific focus on expanded Tregs. We further applied our autoencoder model for cells having TCR sequence passing our quality control, and obtained their expansion probability. The area under the ROC curve was then calculated separately by the different sample types: adjacent normal, blood, and tumor samples (Figure S8C-D). Using optimal Treg expansion threshold in tumor samples for the autoencoder model (Figure S2A), we labeled Tregs predicted as expanded by the model and calculated the fraction of both truly expanded and predicted expanded Tregs out of all the T cells per sample (Figure 6G-I). We used a two-sided Wilcoxon rank-sum test to compare this fraction between patients based on their cancer-neural invasion status.

### Quantification and statistical analysis

The quantitative and statistical analyses are described thoroughly in the relevant sections of the method details. Explicit p-values and sample sizes are embedded in the figures and/or figure legends, and are described throughout the text.

## Supplemental information titles and legends

**Document S1. Figures S1-S8.**

**Table S1. Summary of utilized paired scRNA/TCR-seq datasets,** Related to Figures 1, 4, 6, S1, S3-5, S8.

**Table S2. Model hyperparameters and performance metrics,** Related to Figures 2-3, S2.

**Table S3. Model performance across external independent test datasets,** Related to Figures 4, S3-5.

## Notes

### Competing Interest Statement

The authors have declared no competing interest.

